# Behavioral tagging and the penumbra of learning

**DOI:** 10.1101/072678

**Authors:** Samuel J. Gershman

**Affiliations:** Department of Psychology and Center for Brain Science Harvard University

## Abstract

In noisy, dynamic environments, organisms must distinguish genuine change (e.g., the movement of prey) from noise (e.g., the rustling of leaves). Expectations should be updated only when the organism believes genuine change has occurred. Although individual variables can be highly unreliable, organisms can take advantage of the fact that changes tend to be correlated (e.g., movement of prey will tend to produce changes in both visual and olfactory modalities). Thus, observing a change in one variable provides information about the rate of change for other variables. We call this the *penumbra of learning.* At the neural level, the penumbra of learning may offer an explanation for why strong plasticity in one synapse can rescue weak plasticity at another (synaptic tagging and capture). At the behavioral level, it has been observed that weak learning of one task can be rescued by novelty exposure before or after the learning task. Here, using a simple number prediction task, we provide direct behavioral support for the penumbra of learning in humans, and show that it can be accounted for by a normative computational theory of learning.

## Introduction

Some of the most significant developments in synaptic physiology have sprung from the insight that no synapse is an island: changes in plasticity at one synapse can exert a variety of effects on plasticity at other synapses, such as synaptic scaling [1], heterosynaptic plasticity [2], and (the focus of this paper) synaptic tagging and capture [3, 4, 5]. The synaptic tagging and capture hypothesis holds that weak stimulation at a synapse, while insufficient to produce synthesis of plasticity-related proteins necessary for long-lasting potentiation, sets a molecular tag that allows the synapse to subsequently capture proteins synthesized at another convergent synapse undergoing strong stimulation. Capture is possible in both a “weak-before-strong” scenario (weak stimulation at synapse A followed by strong stimulation at synapse B) and a “strong-before-weak” scenario (strong stimulation at synapse B followed by weak stimulation at synapse A). Tagging and capture also occurs for synapses undergoing depression [6]. Furthermore, strong depression at one synapse can rescue weak potentiation at another synapse, and vice versa (“cross-capture” [6]).

In an effort to organize the functional principles underlying these phenomena, Gershman [7] proposed a normative computational model of synaptic tagging and capture. According to this model, a basic challenge facing any learning system is distinguishing between change in the environment (which should be learned) and noise (which should be ignored). Under the assumption that the rate of change is correlated across different variables, observing a large change for one variable is informative about the global rate of change, which in turn alters learning about other changes—the *penumbra of learning.* Thus, changes that appear insignificant under some circumstances (due to noise) may attain significance when the environment is perceived to be changing quickly. Applied to the synaptic level, this principle explains why strong learning at one synapse can enhance weak learning at another synapse.

One advantage of such an abstract model formulation is that it can be applied at multiple levels; for example, we should be able to observe the penumbra of learning behaviorally. Indeed, “behavioral tagging” experiments demonstrate that exposing an animal to novelty can enhance weak learning on a separate task [8, 9, 10, 11]. One limitation of these experiments is that they are not amenable to a systematic computational analysis, since it is unclear how to quantitatively model the novelty manipulation. In the present paper, we study a simple behavioral task that allows us to test the model’s predictions in humans.

Participants performed a number prediction task [12] in which they had to guess the number a colored button would produce. Each button’s output changed over time according to a Gaussian random walk, such that participants had to continually update their predictions. Trials alternated between red and green buttons Figure 1, allowing us to measure the penumbra of learning in terms of whether a large or small change in prediction for one of the buttons was associated with a large or small change in the other button. By looking at the direction of change (i.e., positive or negative), we can also measure a behavioral analogue of cross-capture: an association between large positive changes on one button with large negative changes in the other button, or vice versa.

**Figure 1:**
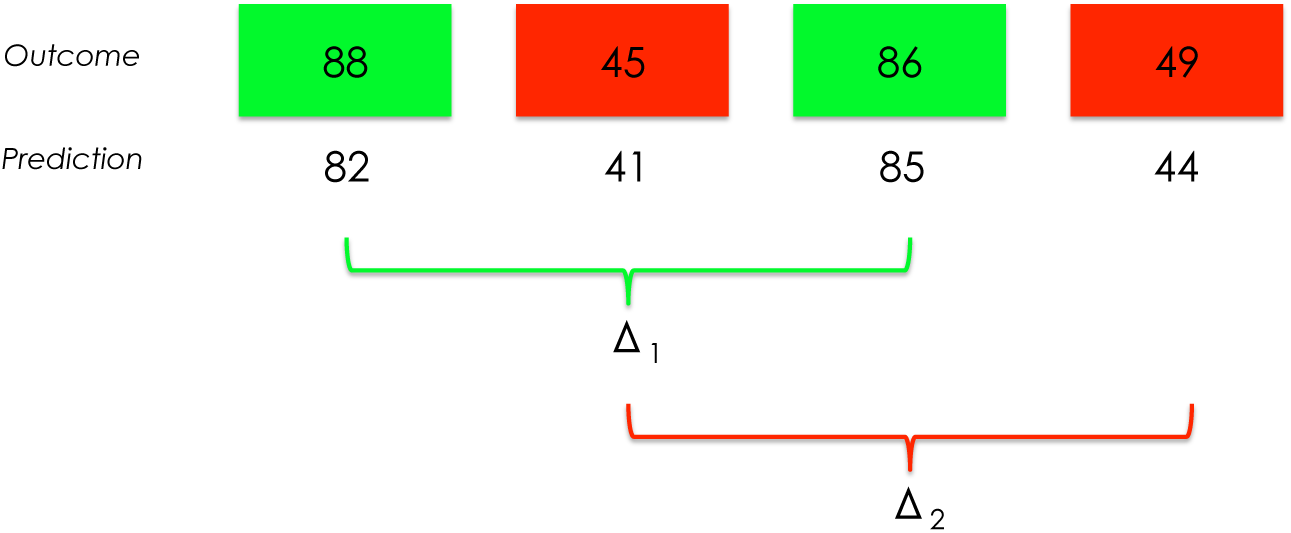
Task and analysis schematic. Participants alternated between making predictions for red and green buttons. After entering a predicted number, participants observed feedback about the true number, drawn from a Gaussian distribution whose mean fluctuated over time according to a Gaussian random walk. Two variables were computed from the sequence of predictions: the difference between contiguous green predictions (Δ_1_) and the difference between contiguous red predictions (Δ_2_). These variables were used to quantify the penumbra of learning, as explained in the main text.

## Results

### Experiment 1

The experimental results are summarized in Figure 2. First, we measured “forward tagging”: the absolute change in prediction on trial *t* (|Δ_2_|) as a function of whether the absolute change in prediction on trial *t* – 1 (|Δ_1_|) is large (higher than average) or small (lower than average). Consistent with the penumbra of learning hypothesis, we observed that |Δ_2_| was larger when |Δ_1_| was large compared to when |Δ_x_| was small [*t*(148) = 6.23, *p* < 0.0001; Figure 2A]. We also observed “backward tagging”: |Δ_X_| was larger when |Δ_2_| was large compared to when | Δ_2_ | was small [*t*(148) = 5.47, *p* < 0.0001; Figure 2B].

**Figure 2:**
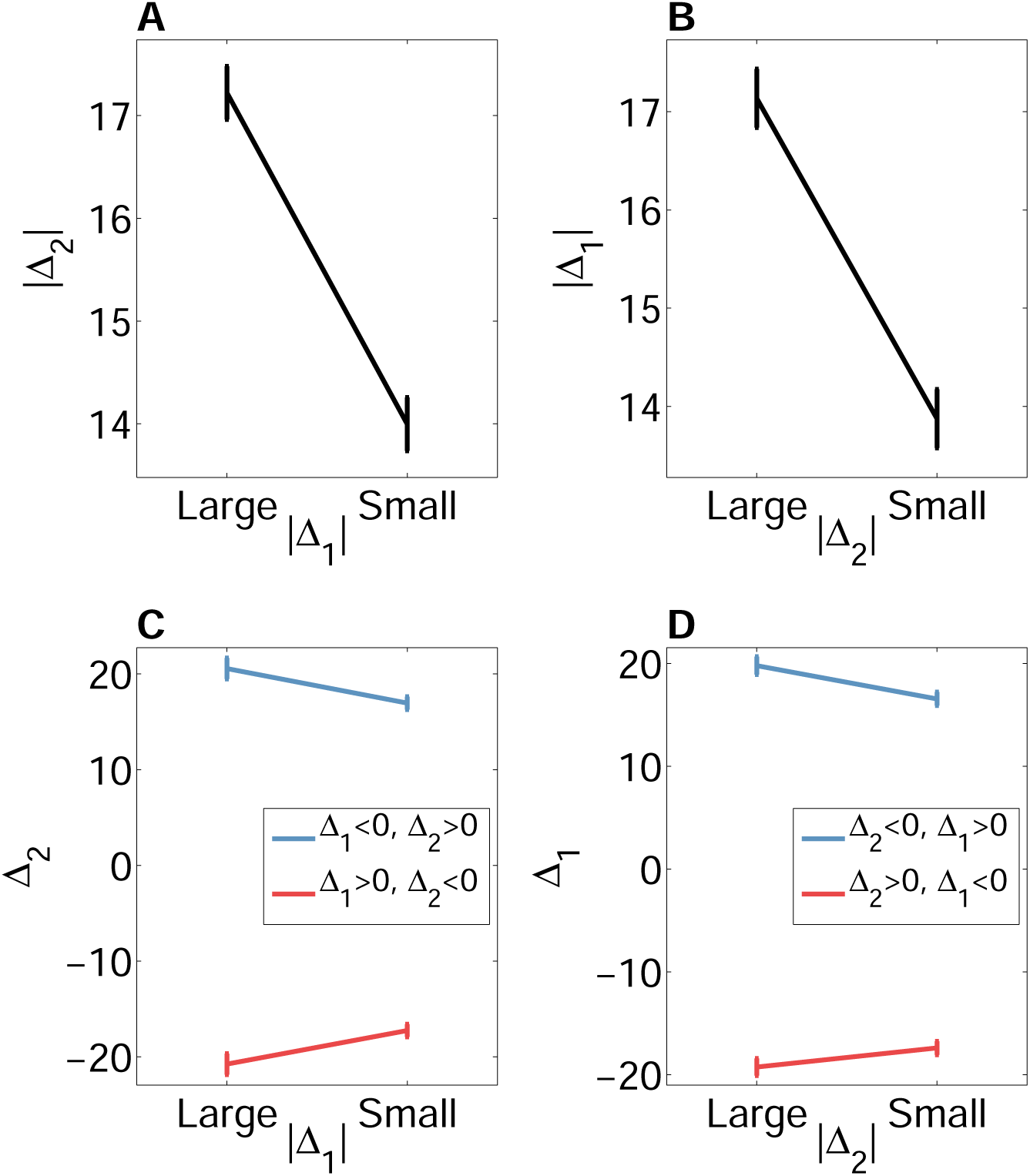
Experiment 1 results. (A) Forward tagging: absolute change in prediction on trial t as a function of the size of change in prediction on trial *t* – 1. The “large” and “small” categories were determined on the basis of a mean split. (B) Backward tagging: absolute change in prediction on trial *t* – 1 as a function of the size of change in prediction on trial *t*. (C) Forward cross-tagging: a negative change on trial *t* – 1 amplifies a positive change on trial *t*. Similarly, a positive change on trial *t* – 1 amplifies a negative change on trial *t*. (D) Backward cross-tagging: a negative change on trial t amplifies a positive change on trial *t* – 1. Similarly, a positive change on trial *t* amplifies a negative change on trial *t* – 1. Error bars show standard error of the mean.

We next examined a behavioral analogue of “cross-capture,” where a change in prediction for one button amplifies change in prediction for the other button, even when the directions of these changes go in opposite directions. Thus, a negative change on trial £ – 1 amplifies a positive change on trial *t* [*t*(117) = 3.92, *p* < 0.001; Figure 2C], and a positive change on trial *t* – 1 amplifies a negative change on trial *t* [*t*(120) = 3.44, *p* < 0.001; Figure 2D]. These results support the claim that it is the magnitude, not direction, of change that determines the strength of the penumbra. Consistent with this claim, we found no significant correlation between predictions on trial *t* and trial *t–* 1 (median Pearson correlation coefficient: 0.06, *p* = 0.15, signed rank test).

These empirical results are well captured by the normative computational model described in the Methods section Figure 3, which was previously used to explain a variety of synaptic tagging and capture phenomena [7]. The key principle is that participants are simultaneously estimating both the number and the diffusion variance (or volatility) *q*. Change on one button is, according to the model, informative about the rate of change for all buttons. Formalized mathematically, without any parameter fitting, this principle is sufficient to reproduce forward/backward tagging and forward/backward cross-tagging.

**Figure 3:**
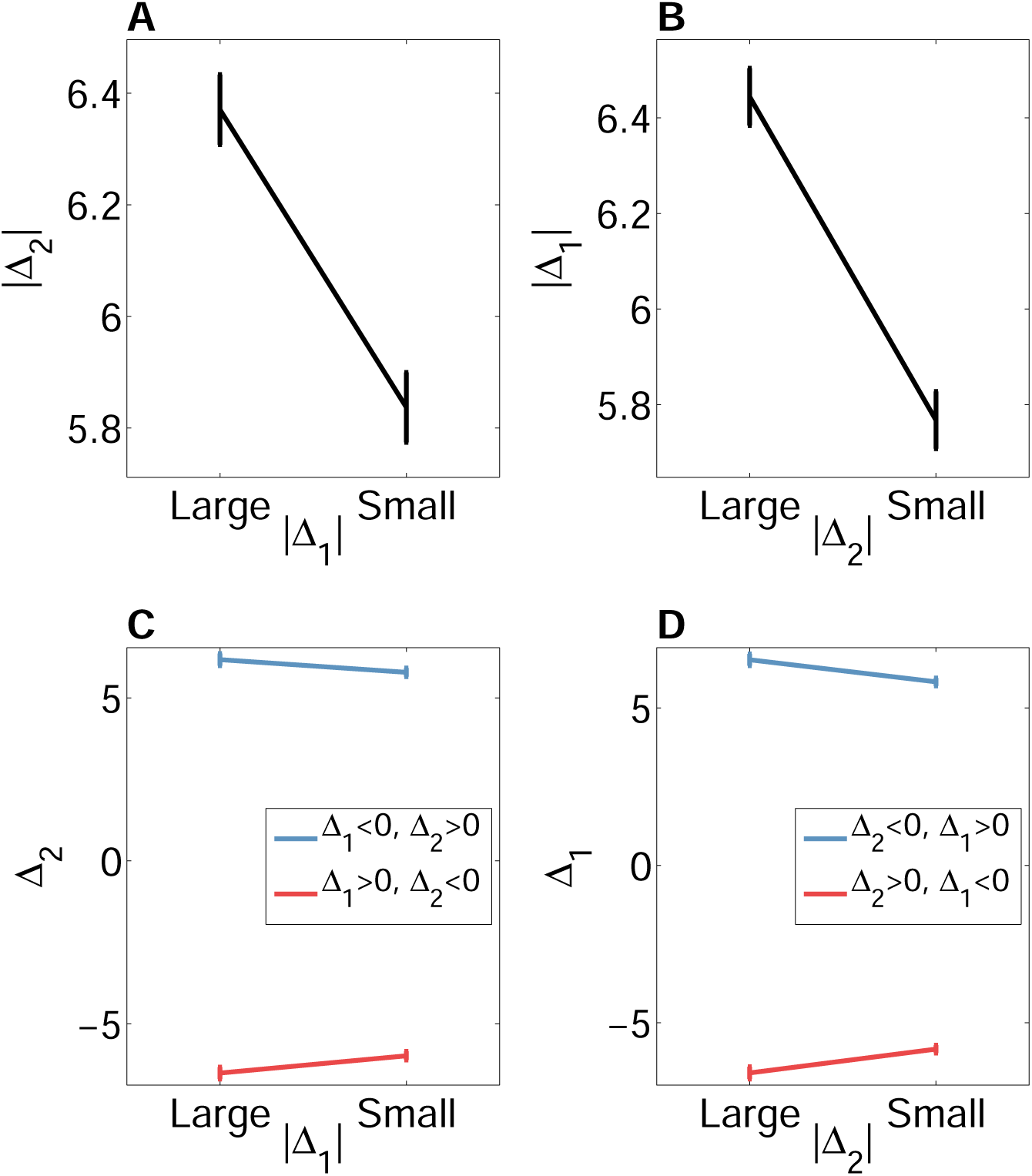
Experiment 1 model simulations. (A) Forward tagging. (B) Backward tagging. (C) Forward cross-tagging. (D) Backward cross-tagging. Error bars show standard error of the mean.

### Experiment 2

In Experiment 1, the diffusion variance was the same across buttons, and we exploited natural variability in prediction errors to test the model’s predictions. In Experiment 2, we sought to directly manipulate the size of prediction errors for button 1 while keeping the prediction errors for button 2 constant. A separate set of participants (*N* = 46) performed the same task as in Experiment 1, with diffusion variance for button 1 (*q*_2_ set to 225 and the diffusion variance for button 2 (*q*_2_) set to 25. We refer to this new condition as “high volatility” for button 1, compared to the “low volatility” condition studied in the subset of participants in Experiment 1 (*N* = 53) for whom both *q*_1_ and *q*_2_ were set to 25 (see Methods for details). Crucially, the diffusion variance for button 2 was the same across the high volatility and low volatility conditions.

As shown in Figure 4, we found that the absolute change in prediction for button 2 (|Δ_2_|) was significantly greater in the high volatility condition compared to the low volatility condition [*t*(97) = 2.19, *p* < 0.05]. This finding supports the model prediction that high volatility for one button will induce greater plasticity of learning for the other button (whose volatility was fixed). Experiment 2 thus complements the correlational results of Experiment 1 with a causal experimental manipulation.

**Figure 4:**
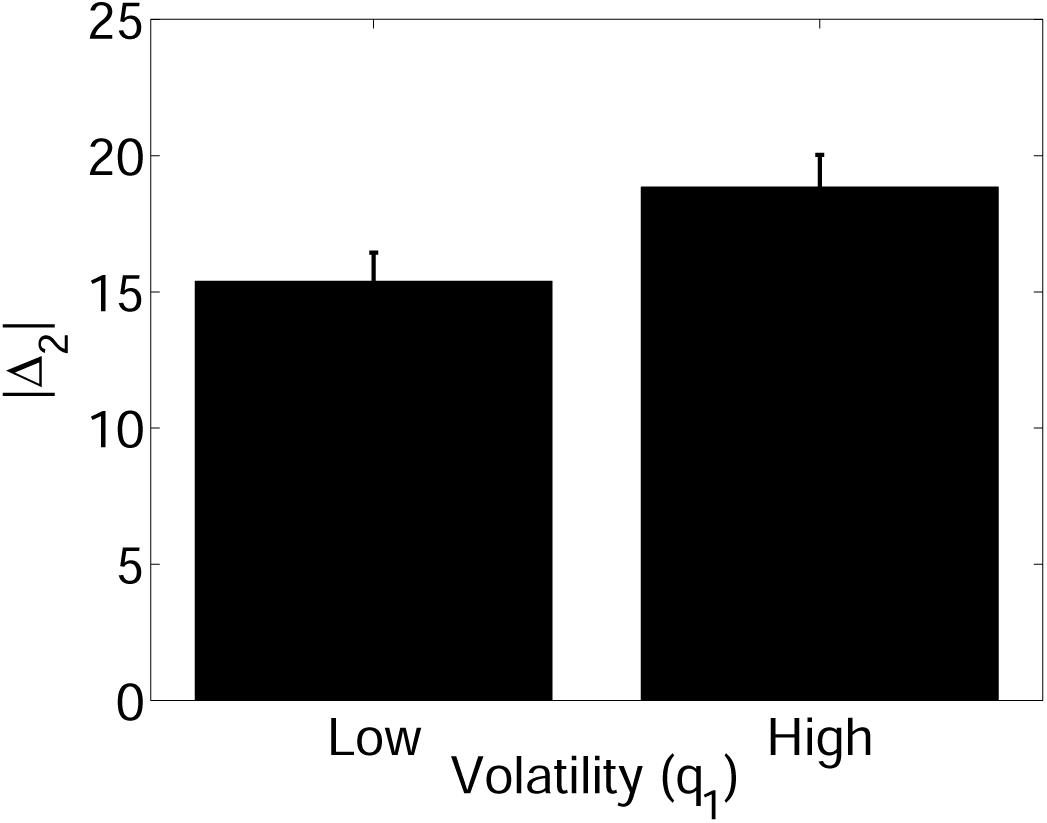
Experiment 2 results. Absolute change in prediction for button 2 (|Δ_2_|) as a function of volatility for button 1 (Low volatility: *q*_1_ = 25; High volatility: *q*_1_ = 225.). Error bars show standard error of the mean.

## Discussion

We have shown a form of behavioral tagging in humans that recapitulates analogues of several important synaptic tagging and capture phenomena. First, we found that a large change in predictions for one stimulus amplified learning about another stimulus. This “penumbra of learning” operated both forward and backward in time. Second, we found that a large positive change for one stimulus amplified negative changes for the other stimulus, and vice versa, indicating that the penumbra depends on the magnitude (rather than direction) of change. Third, we found that increasing the volatility (and hence the average change in predictions) for one stimulus amplified learning about the other stimulus, indicating that the penumbra of learning can be exogenously manipulated.

Our paradigm made it possible to test a computational account of synaptic tagging and capture [7], applied with minimal modifications to the behavioral task. This model formalizes the penumbra of learning in terms of inference about the volatility of the environment. Under the assumption that stimulus dynamics are coupled via an unknown volatility parameter, perceiving a change in one stimulus increases the inferred volatility and thereby amplifies learning about other stimuli. The notion that humans and animals adapt their learning rate based on inferred volatility has become an important facet of modern learning theory [13, 14, 15, 16], with roots in earlier models of classical conditioning [17]. The penumbra of learning model extends this logic to multidmensional environments. Although originally applied to synaptic plasticity phenomena [7], we have shown that the model can accurately predict the behavioral tagging phenomena reported in this paper, lending credence to the idea that synaptic and behavioral learning reflect common principles. However, we still lack physiological evidence for a direct connection.

Our results pave the way for several future directions of inquiry. First, it has been demonstrated in animals that behavioral tagging is dopamine-dependent [8, 18, 10], suggesting that systemically depleting dopamine in humans should eliminate the penumbra of learning. Behavioral tagging is also dependent on β-adrenergic function [18], which could also be systemically manipulated. The role of β-adrenergic function is particularly intriguing given the role of noradrenergic neurons in adaptation of learning [19, 20]. Pupil diameter, tightly linked to firing of noradrenergic neurons [20], tracks changes in learning rate [15], and could be easily measured in our task.

## Methods

### Participants

For Experiment 1, 149 participants were recruited online through the Amazon Mechanical Turk web service and paid for their participation. For Experiment 2, 46 participants were recruited. The experiments were approved by the Harvard Internal Review Board.

### Procedure

Participants completed 100 trials of a number prediction task. On each trial, participants were shown a red or green button (on alternating trials) and had to guess a number between 1 and 100. Participants then received feedback. The numbers generated by the buttons evolved over time according to a Gaussian random walk:

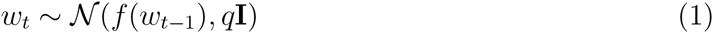

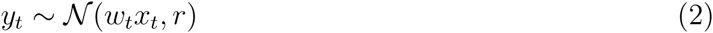

where *y*_*t*_ is the number observed on trial *t*, *w*_*t*_ ∈ *ℝ*^2^ is the vector of mean values, I is the 2 × 2 identity matrix, *x*_*t*_ ∈ ℝ^2^ is a vector representing the button presented on trial *t* (*x*_*tk*_ = 1 if button *k* was present on trial *t*, and 0 otherwise), *q* is the diffusion variance of the random walk, and *r* is the noise variance of the emission distribution. The function *f*(*w*) specifies the deterministic component of the dynamics; here we assume *f*(*w*) = *w*, but in the next section we discuss different assumptions about *f*(*w*). To explore the parameter space in Experiment 1, 53 participants completed the task with *q* = 25 and *r* = 25, 50 with *q* = 1 and *r* = 25, and 46 with *q* = 25 and *r* = 100. However, the experimental results were nearly indistinguishable across conditions, so in the remainder of the paper we report results pooled across conditions. Experiment 2 used an identical procedure, with the only difference that the two buttons had different diffusion variances: *q*_1_ = 225 and *q*_2_ = 25.

### Computational model

Here we summarize the computational model first presented in [7]; we refer the reader to that paper for details. Our approach is to impute to the learner a set of probabilistic assumptions about the environment (the *generative process*), and then formalize the learning problem as inference over the latent causes that gave rise to observed data under the generative process. Thus, we posit that the learner is executing a reasonable accurate approximation of probabilistic inference. In [7], the learner was conceptualized as a set of synapses; in this paper, we apply essentially the same model at the organismic level, only changing some of the parameters to match our experimental paradigm.

Formally, the computational problem facing the learner is to infer both the dynamic mean *w*_*t*_ and the diffusion variance *q* after observing data (x_1:*t*_, y_1:*t*_), where the subscript 1: *t* denotes the collection of variables on trials 1 to *t*. The optimal statistical inference is stipulated by Bayes’ rule:

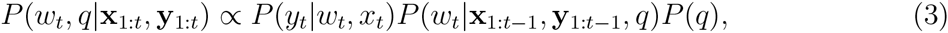

We approximate the distribution over the diffusion variance *q* with a discrete set of points, {*q*^1^, ⋯, *q*^*K*^}. Specifically, the diffusion variance is drawn according to *P*(*q* = *q*^*k*^) ∝ exp{*ak*^-*b*^} where *a* and *b* are constants and *q*^*k*^ ranges over *K* equally spaced values between 1 and 30 (note that the *k* superscript is an index, not a power). This distribution embodies the assumption that a slow rate of change is *a priori* more probable. For all our simulations, we used *K* = 50, *a* = 8 and *b* = 5, but the results are not sensitive to small variations in these values.

When *q* is known, the posterior over *w*_*t*_ can be approximated as a Gaussian using the extended Kalman filter [21], leading to the following updates for the mean and variance:

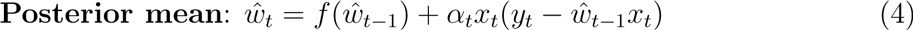

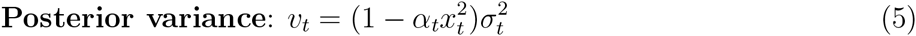

where 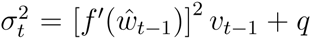 is the predictive variance after application of the transition function 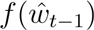 but before observing (*x*_*t*_,*y*_*t*_), and *fʹ*(*w*) is the derivative of the transition function.

The learning rate α*_t_* is given by 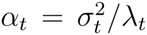, where 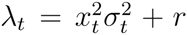 is the variance of the posterior marginal likelihood:

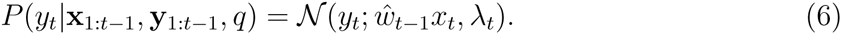

Intuitively, λ*_t_* expresses the overall unpredictability (or noisiness) of the data; this unpredictability grows monotonically with the noise variance. Learning will be faster to the extent that the prediction uncertainty is high (large 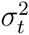) relative to the overall unpredictability (low *At*).

With *q* unknown, the posterior marginal over λ*t* is a mixture of Gaussians:

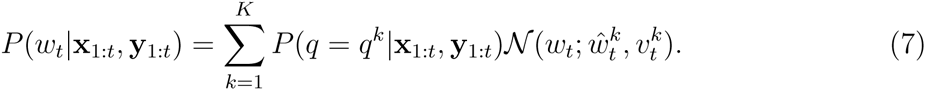

where we have indexed ŵ and *v* by *k* to indicate their implicit dependence on *q* = *q*^*k*^. To obtain *P*(*q* = *q*^*k*^|**x**_1:_*_t_*,**y**_1:_*_t_*), we apply the update equations 4 and 5 for each *q*^*k*^ in parallel, and use Bayes’ rule to compute the posterior over *q*:

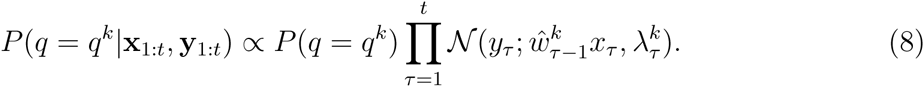

In [7], a nonlinear transition function was specified that captures two properties of synaptic plasticity: (1) asymptotic decay to 0, and (2) slower decay for stronger associations. The following functional form was used:

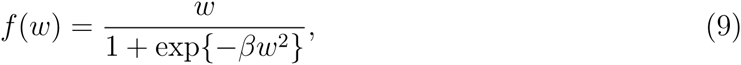

where β ≥ 0 is a parameter that governs the nonlinear relationship between *w* and the decay rate; in particular, smaller absolute values of *w* decay more rapidly than larger values. Following [7], we used β = 10. Note that in the *true* generative process (unknown to participants), *f* (*w*) was the identity function. We employ the nonlinear transition function for consistency with prior work, but our results do not depend on this assumption.

In the simulations, all parameters were the same as in the original report [7], except that we set *r* equal to its true experimental value (25 or 100 depending on the experimental condition). Participants did not know this true value, but we chose it for expedience; our results are robust to large variations in this parameter value.

## Acknowledgments

This work was supported in part by the Center for Brains, Minds and Machines (CBMM), funded by NSF STC award CCF-1231216.

